# A series of dual-reporter vectors for ratiometric analysis of protein abundance in plants

**DOI:** 10.1101/2020.02.10.939363

**Authors:** Aashima Khosla, Cecilia Rodriguez-Furlan, Suraj Kapoor, Jaimie M. Van Norman, David C. Nelson

**Affiliations:** Department of Botany and Plant Sciences, University of California, Riverside, CA 92521 USA; Department of Genetics, University of Georgia, Athens, GA 30602 USA

## Abstract

Ratiometric reporter systems enable comparisons of the abundance of a protein of interest, or “target,” relative to a reference protein. Both proteins are encoded on a single transcript but are separated during translation. This arrangement bypasses the potential for discordant expression that can arise when the target and reference proteins are encoded by separate genes. We generated a set of 18 Gateway-compatible vectors termed pRATIO that combine a variety of promoters, fluorescent and bioluminescent reporters, and 2A “self-cleaving” peptides. These constructs are easily modified to produce additional combinations or introduce new reporter proteins. We found that mScarlet-I provides the best signal-to-noise ratio among several fluorescent reporter proteins during transient expression experiments in *Nicotiana benthamiana*. Firefly and Gaussia luciferase also produce high signal-to-noise in *N. benthamiana*. As proof of concept, we used this system to investigate whether degradation of the receptor KAI2 after karrikin treatment is influenced by its subcellular localization. KAI2 is normally found in the cytoplasm and the nucleus of plant cells. In *N. benthamiana*, karrikin-induced degradation of KAI2 was only observed when it was retained in the nucleus. These vectors are tools to easily monitor *in vivo* the abundance of a protein that is transiently expressed in plants, and will be particularly useful for investigating protein turnover in response to different stimuli.

## INTRODUCTION

Dynamic monitoring of protein abundance *in vivo* requires an easily detectable reporter system. Translational fusions of fluorescent or bioluminescent proteins to a protein of interest, here referred to as a “target,” are commonly used for this purpose (Bronstein et al., 1994; Wood, 1995; Genové et al., 2005). Stable transformation of a host organism with the target-encoding construct is typically carried out to produce a replicable and relatively homogeneous reporter line, although genetic instability or gene silencing may occur over subsequent generations (Vaucheret et al., 1998). The local genetic context (e.g. chromatin state or nearby enhancer elements) can influence the expression of a transgene. Therefore, isolation of multiple homozygous transgenic lines is typically required to identify one with consistent and readily detectable expression of the target. For many experiments, the cost and time involved in developing many transgenic reporter lines can discourage rapid progress. This can be resolved with transient transformation and expression of a reporter construct, which in plant biology research is often carried out in protoplasts or *Nicotiana benthamiana* (hereafter referred to as tobacco) leaves (Yang et al., 2000; Wroblewski et al., 2005). In transient expression experiments, a second reporter protein that can function as a reference is useful to normalize for differences in transformation efficiency or transgene expression across samples.

There are several ways to achieve a dual reporter system. Perhaps the most commonly used approach is co-transformation of separate target- and reference-encoding plasmids (Larrieu et al., 2015; De Sutter et al., 2005). This does not guarantee that both constructs enter each cell, or that they do so with consistent proportions. Differences in the size of each plasmid may also impact their relative transformation efficiencies. An improvement on this method is to encode the target and reference protein on the same plasmid with each regulated by its own promoter and transcriptional termination sequence (Moyle et al., 2017; Koo et al., 2007). This can work well, but potential problems include the increased plasmid size, the relative activity of the two promoters if they are different, and recombination between promoters or terminators if they are identical.

Furthermore, the order in which the genes are expressed in the vector can influence their expression levels (Halpin, 2005).

An attractive third option is to encode the target and reference protein on the same transcript. This avoids variation in the relative expression of the target and reference genes that may occur in different cell types or environmental conditions, as transcription of both genes will be affected equally. Multicistronic gene expression can be achieved in eukaryotes through incorporation of an internal ribosome entry site (IRES) or a sequence encoding a 2A “self-cleaving” peptide between the target and reference genes. IRES sequences produce a secondary structure in the mRNA that enables translation to occur downstream (Urwin et al., 2000). The efficiency of translation for proteins encoded upstream and downstream of the IRES can vary widely depending on the IRES sequence selected (Urwin et al., 2002). Studies comparing the expression levels of two cDNA sequences separated by an IRES have shown that genes cloned downstream of the IRES were expressed at significantly lower levels (10 – 50% of the upstream gene) (Mizuguchi et al., 2000). A second drawback of IRES sequences is that they are somewhat large, typically ∼500 to 600 bp. Because of this, 2A peptides have become a commonly used alternative to produce multicistronic expression in eukaryotes.

The 2A peptide from foot-and-mouth disease virus (FMDV, “F2A”) and several 2A-like sequences are able to disrupt normal translation, causing a protein encoded downstream of 2A to be translated separately from a protein encoded upstream (Halpin et al., 1999; Ralley et al., 2004; Luke et al., 2015). This “ribosome skipping”, “stop-go”, or “self-cleavage” effect occurs when the ribosome fails to create a glycyl-prolyl bond at the end of the 2A peptide but then continues translation (Atkins et al., 2007; Doronina et al., 2008). The first protein retains the majority of the 2A peptide as a C-terminal fusion, while the proline residue becomes the N-terminus of the second protein (Donnelly et al., 2001c, 2001a; Luke and Ryan, 2018). Cleavage is thought to be caused by interaction of the nascent 2A peptide with the ribosomal exit tunnel. Indeed the length of the 2A peptide impacts its cleavage efficiency. A minimum of 13 amino acids of F2A are required for cleavage, but longer versions are more effective. Including residues from the 1D capsid peptide encoded upstream of 2A in FMDV can further increase the cleavage efficiency of a 2A sequence (Donnelly et al., 2001a; Minskaia et al., 2013). While longer versions have been shown to produce the most efficient cleavage, in some cases the ‘remnant’ 2A residues appended to the C-terminus of a processed protein may hinder its activity (François et al., 2004; Randall, 2004; Samalova et al., 2006). Removal of the extraneous 2A residues using endogenous proteases has been attempted in plant (François et al., 2004) and mammalian systems (Fang et al., 2005). Conversely, when shorter 2A sequences are used the C-terminal sequence of the upstream protein can impact cleavage efficiency (Minskaia et al., 2013). For shorter 2As, cleavage efficiency has been improved by insertion of a flexible Gly-Ser-Gly or Ser-Gly-Ser-Gly spacer sequence between the upstream protein and the 2A sequence (Fang et al., 2005; Lorens et al., 2004; Szymczak et al., 2004a; Provost et al., 2007).

Several studies have used 2A peptides in dual reporter systems in plants. For example, Wend *et al*. developed a degradation-based biosensor to study auxin dynamics in transient expression systems (Wend et al., 2013). The chemiluminescent sensor is composed of two components: a Aux/IAA degron fused to firefly luciferase (LUC) as a target, and Renilla luciferase as a reference. Both components are linked by a 23 aa F2A peptide. In the presence of auxin and the co-receptor F-box protein TIR1, the target is degraded. Samodelov *et al*. used a similar degradation-based sensor, termed StrigoQuant, to monitor strigolactone signaling in plant protoplasts (Samodelov et al., 2016). This construct expressed SUPPRESSOR OF MORE AXILLARY GROWTH2-LIKE6 (SMXL6) fused to LUC as a target, and Renilla luciferase as a reference. Separation of the target and reference proteins was achieved by the same 23 aa F2A peptide as Wend et al. The SMXL6-LUC target is degraded after strigolactone perception. Samalova *et al*. utilized a fluorescent reporter to study plant membrane trafficking in both transiently and stably transformed systems (Samalova et al., 2006). A 20 aa F2A peptide was used to co-express a trafficked fluorescent protein marker in fixed stoichiometry with a reference fluorescent protein localized to a different cellular compartment.

We are interested in developing a similar ratiometric system to report on signaling activity in the karrikin pathway in plants. Karrikins (KARs) are a class of butenolide compounds found in smoke that can stimulate seed germination and enhance the photomorphogenic growth of *Arabidopsis thaliana* seedlings (Flematti et al., 2004; Nelson et al., 2009, 2010, 2012). KAR responses in plants require the a/b-hydrolase protein KARRIKIN INSENSITIVE2 (KAI2)/HYPOSENSITIVE TO LIGHT(HTL) (Waters et al., 2012; Sun and Ni, 2011). *KAI2* has roles in germination, hypocotyl elongation, drought tolerance, root skewing, root hair development, and symbiotic interactions with arbuscular mycorrhizal fungi (Gutjahr et al., 2015; Li et al., 2017; Villaécija-Aguilar et al., 2019; Swarbreck et al., 2019). In addition to mediating KAR responses, it is thought that KAI2 recognizes an unknown, endogenous signal known as KAI2 ligand (KL) (Conn and Nelson, 2015). If so, KARs might be natural analogs of KL, to which some fire-following species have become particularly attuned.

KAI2 works with the F-box protein MORE AXILLARY GROWTH2 (MAX2) to mediate KAR responses, likely through polyubiquitination and degradation of SUPPRESSOR OF MAX2 1 (SMAX1) and SMAX1-LIKE2 (SMXL2) (Nelson et al., 2011; Stanga et al., 2013, 2016). KAR treatment causes degradation of KAI2 protein over the course of several hours, putatively as a form of negative feedback regulation (Waters et al., 2015). Proteolysis of KAI2 occurs independently of MAX2 through a mechanism that is currently unknown. Substitution of Ser95, one of the catalytic triad residues, with alanine renders KAI2 non-functional and also prevents its degradation in the presence of KAR_2_ (Waters et al., 2015). Potentially, KAI2 degradation could be used as the basis of an *in vivo* reporter for its activation. Such a bioassay could be useful in attempts to identify KL through fractionation of small molecule extracts from plants.

This led us to develop a series of Gateway-compatible, plant transformation vectors for ratiometric detection of a protein of interest in transient expression assays. We tested the cleavage efficiency of two versions of the foot-and-mouth disease virus (FMDV) 2A peptide. We compared the signal-to-noise ratio of several fluorescent and bioluminescent reporters transiently expressed in *Nicotiana benthamiana* to identify those with the largest potential dynamic range. Finally, as proof-of-concept, we used the ratiometric system to investigate KAR-activated proteolysis of KAI2.

## RESULTS

### Design of pRATIO vectors

We constructed a series of 18 Gateway-compatible binary vectors named pRATIO that encode multicistronic ratiometric reporters (Figure 1). A gene of interest can be transferred readily from an entry clone into the destination vector through an LR Gateway reaction (Invitrogen). The target is composed of a gene of interest that has an in-frame, C-terminal fusion to a fluorescent or bioluminescent reporter gene. This is followed by a 2A peptide-encoding sequence and a second fluorescent or bioluminescent reporter gene that serves as a reference. After the 2A peptide interrupts translation the ribosome may fall off instead of resuming translation of the next coding sequence; typically this results in a higher molar ratio of the first protein product vs. the second (Donnelly et al., 2001b; Liu et al., 2017). Therefore, to maximize target signal, we chose to encode the target protein first. Expression of the multicistronic transcript is controlled by a single promoter and nopaline synthase terminator (T_nos_). We selected the *35Sp* from cauliflower mosaic virus, which is commonly used to drive strong expression of transgenes in plants. However, *35Sp* is not equally expressed across all tissue types and can be prone to silencing (Elmayan and Vaucheret, 1996). To achieve more uniform expression of the ratiometric construct, several pRATIO vectors carry the *UBIQUITIN 10* promoter (*UBQ10p*) from *Arabidopsis thaliana. UBQ10p* works well for transient expression in *Arabidopsis* and tobacco tissues, and is equally useful for generating stable transgenic lines (Grefen et al., 2010). Some pRATIO vectors include a nuclear localization sequence (NLS) from the SV40 large T antigen that is translationally fused to the N-terminus of the target. In some cases, the presence of an NLS can facilitate the detection of weak fluorescent reporter signals by concentrating the signal in the nucleus (Takada and Jürgens, 2007).

**Figure 1.**
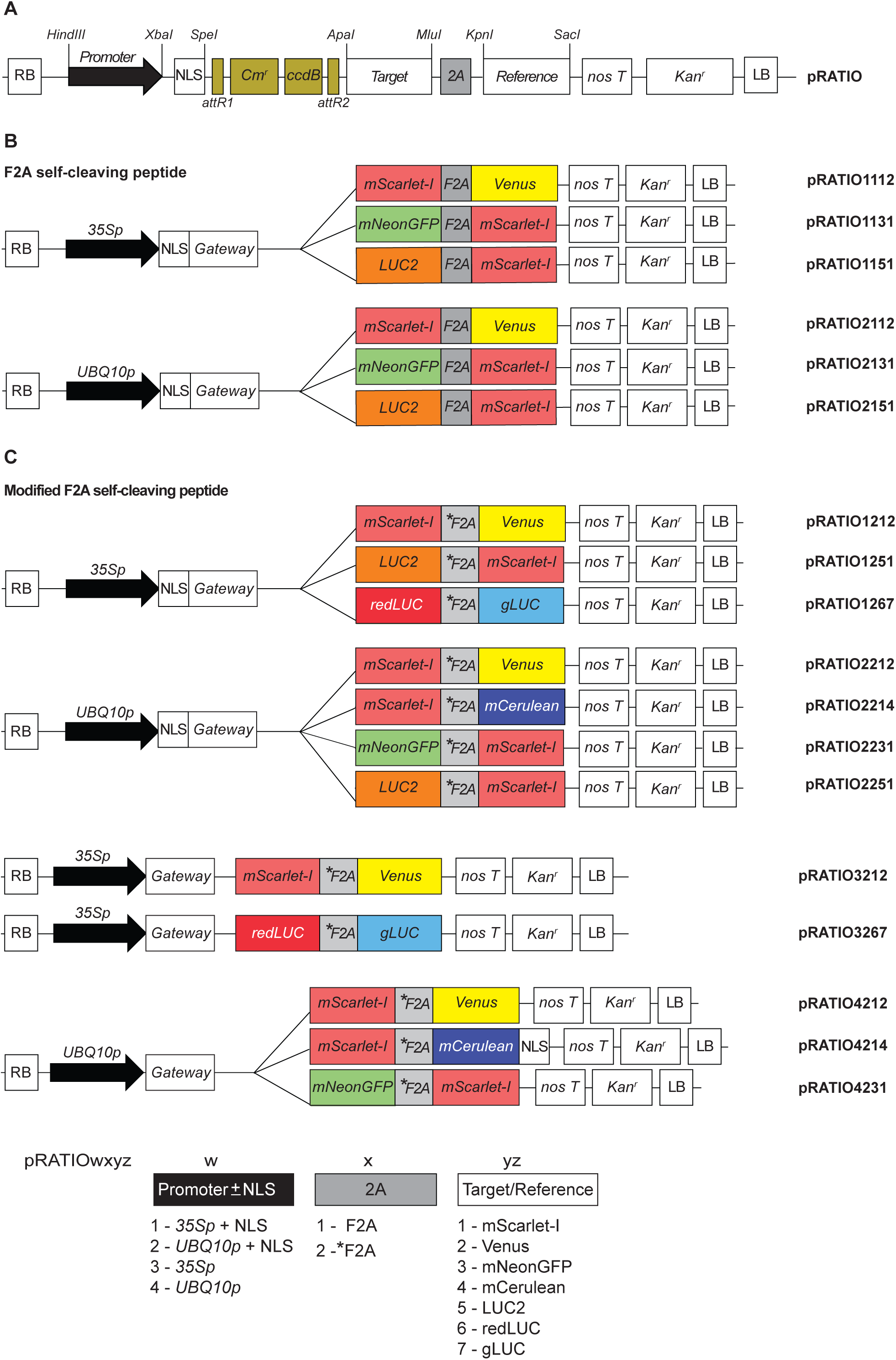
Schematic of the pRATIO vector series. **(A)** General structure of pRATIO vector. Expression is either driven by *CaMV35S* (1000/3000 series) or *UBQ10* promoter (2000/4000 series). Unique restriction sites flank the promoter, NLS, ccdB cassette, target protein, 2A sequence, and the reference protein. The expression cassettes are in pGWB401 or pGWB402 backbones. T-DNA selection is kanamycin resistance. **(B, C)** pRATIO incorporating either **(B)** F2A self-cleaving peptide or **(C)** modified F2A (*F2A) protein. LUC2, firefly luciferase; redLUC, red firefly luciferase; gLUC, Gaussia Dura luciferase; RB, right border; LR, left border; Cmr, chloramphenicol-resistance marker (chloramphenicol acetyl transferase) used for selection in bacteria; ccdB, negative selection marker used in the bacteria; nosT, NOS terminator to stop the transcription. The GenBank accession numbers are listed in Table S5.

The pRATIO vectors are designed to allow for independent exchange or dropout of any vector element using unique restriction endonuclease sites (Figure 1A). This can be accomplished through classical restriction enzyme-mediated subcloning techniques. Alternatively, a digested pRATIO vector and an insert fragment that is bordered by 15-bp sequences that match the vector ends can be assembled seamlessly with a commercially available enzyme mix such as NEBuilder (New England Biolabs).

### 2A-mediated cleavage of transiently expressed reporters in tobacco

The target and reference proteins are intended to separate during translation due to the action of an intervening FMDV 2A (F2A) peptide. The F2A peptide itself is 19 aa long and is sufficient for some degree of cleavage. However, longer versions of F2A that include portions of the 1D capsid protein encoded upstream in FMDV typically produce higher levels of cleavage. N-terminal extension of 2A with 5 aa of 1D improves cleavage, but extension with 14, 21, and 39 aa of 1D produces complete cleavage and an equal stoichiometry of the upstream and downstream translation products (Donnelly et al., 2001a; Ryan et al., 1991).

The pRATIO1100 and pRATIO2100 series use a 30 aa version of F2A (11 aa of 1D plus 2A) while the pRATIO1200, pRATIO2200, pRATIO3200, and pRATIO4200 series use a 40 aa F2A sequence (21 aa of 1D plus 2A). We further modified the 40 aa F2A at its N-terminus to include a 9 aa LP4 linker peptide and a flexible Gly-Ser-Gly linker, producing *F2A (Figure 2A). This strategy is based upon a hybrid linker that fuses LP4, the fourth linker peptide of a polyprotein precursor found in *Impatiens balsamina* seed, to a 20 aa F2A peptide (Tailor et al., 1997). LP4 is post-translationally cleaved after the first or second aa, enabling removal of almost the entire linker from the N-terminal protein (François et al., 2002, 2004). Inclusion of a Gly-Ser-Gly linker at the N-terminal end of a 2A peptide can improve cleavage efficiency (Szymczak et al., 2004b; Holst et al., 2006; Kim et al., 2011; Chng et al., 2015).

**Figure 2.**
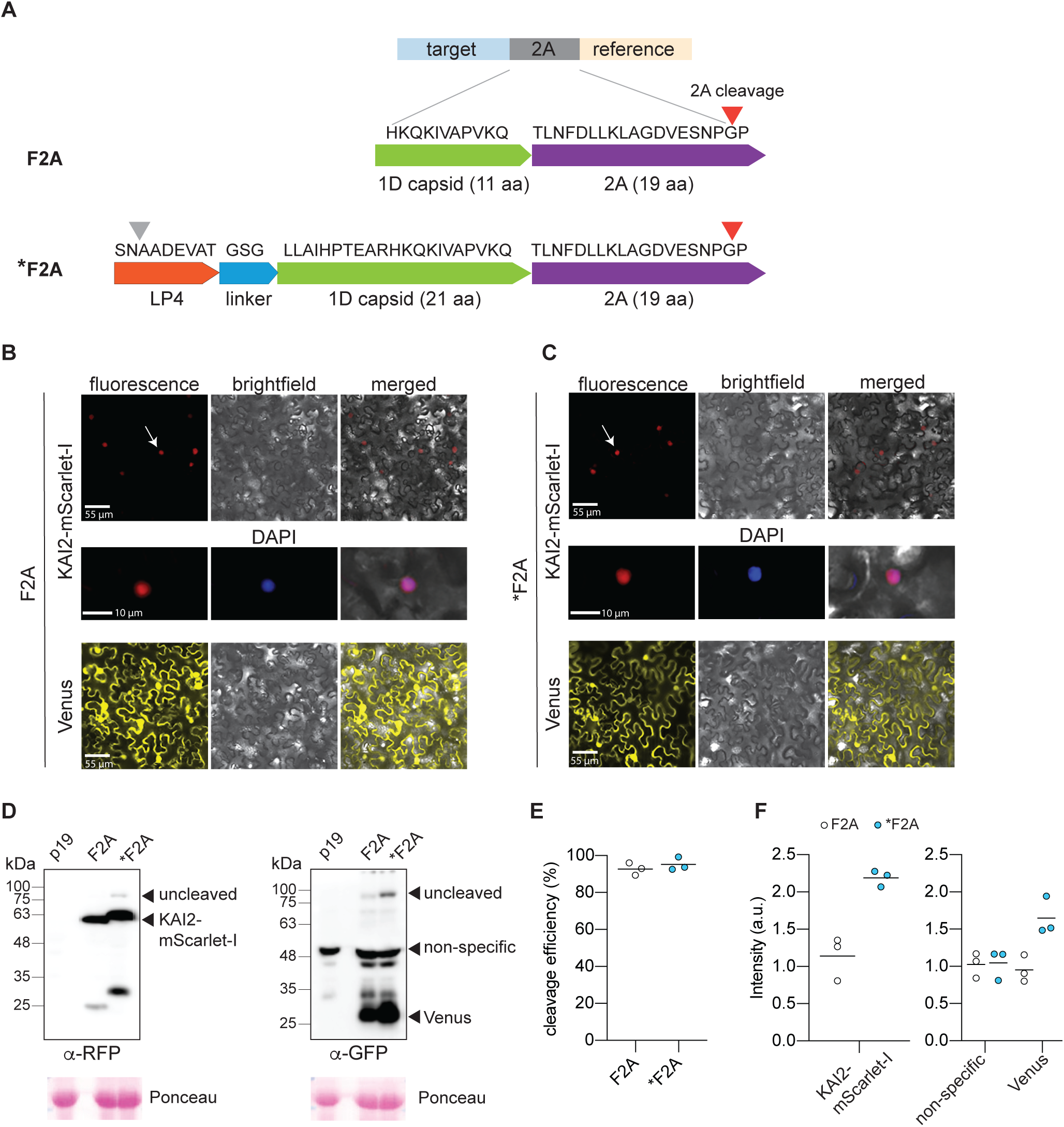
Both F2A and *F2A peptides allow the efficient production of two independent polypeptides in tobacco. **(A)** Amino acid sequences of the two 2As used, foot and mouth disease virus 2A (F2A) and a longer F2A sequence (*F2A). LP4 plant peptide (-SNAADEVAT-) and a GSG linker was added to the N-terminus of *F2A to improve the cleavage efficiency. The site of protease cleavage and 2A mediated cleavage is indicated by green and red arrows, respectively. **(B, C)** Patterns of localization of mScarlet-I and Venus in *N*.*benthamiana* epidermal cells expressing **(B)** pRATIO1112-KAI2 and **(C)** pRATIO1212-KAI2. Venus (yellow) localizes throughout the cells whereas mScarlet-I (red): NLS-KAI2-mScarlet-I remains tightly restricted to the nucleus as expected if they are produced as two independent polypeptides, suggesting that both 2A peptides are correctly split up. Arrows indicate nuclear localization. **(D)** Western blot analysis of cleavage efficiency of two types of 2A self-cleaving peptides in the tobacco leaf epidermal cells. The cleavage efficiency was assessed using RFP antibody to detect cleaved KAI2-mScarlet-I and the uncleaved KAI2-mScarlet-I-2A-Venus, and GFP antibody to detect cleaved Venus and uncleaved protein. Leaf transformed with p19 served as the control. The Ponceau membrane staining of the most intense band at 55 kDa (presumably Rubisco) was used as a loading control. **(E)** Quantitation of cleavage efficiency of 2A peptides in tobacco. Cleavage efficiency = cleaved form/(cleaved form+uncleaved form).The amount of each form was estimated from its band intensity on the Western blot measured by Image Studio Light (LI-COR). Bar indicates mean, n = 3. **(F)**Comparison of the amount of cleaved proteins between pRATIO1112 (F2A) and pRATIO1212 (*F2A). The amount is estimated as depicted in **(E)**. Bar indicates mean, n = 3.

To compare the effectiveness of F2A and *F2A, we cloned an *Arabidopsis KAI2* cDNA into pRATIO1112 and pRATIO1212. We then transiently expressed the constructs in tobacco leaves via *Agrobacterium tumefaciens*-mediated transformation. These reporter systems should produce a NLS-KAI2-mScarlet-I target and a Venus reference protein. Fluorescence microscopy of leaf epidermal cells indicated nuclear localization of mScarlet-I (Figure 2B,C). In contrast, Venus was found in both the cytoplasm and nucleus, as expected for an untargeted monomeric fluorescent protein (FP). These observations were consistent with successful separation of the target and reference proteins.

We further examined the cleavage efficiency of F2A and *F2A through Western blot analysis of total proteins extracted from transiently transformed tobacco leaves. Leaves transformed with p19 alone, which suppresses gene silencing during transient expression, were used as a negative control. We observed very little uncleaved protein (∼87 kDa) compared to NLS-KAI2-mScarlet-I (∼60 kDa) and Venus (27 kDa) in leaves transformed with pRATIO1112-KAI2 and pRATIO1212-KAI2 (Figure 2D). The two versions of 2A peptide performed similarly well; based on the anti-Venus blot we estimated cleavage efficiencies of 92% for F2A and 95% for *F2A (Figure 2E).

The LP4 linker peptide was included to further improve cleavage efficiency through post-translational processing and also to remove the F2A peptide from the C-terminus of the target. We noted that KAI2-mScarlet-I migrated at a slightly higher molecular weight when *F2A was used compared to F2A (Figure 2D). This suggested that cleavage of LP4 might not be occurring as anticipated. Therefore, we probed both samples with a monoclonal antibody against 2A peptide. We found that KAI2-mScarlet-I had retained its 2A peptide in pRATIO1212-KAI2 samples, indicating the LP4 linker component of *F2A was not effective (Supplemental Figure 2).

All considered, *F2A did not appear to offer a clear advantage over F2A; the cleavage efficiencies of these two peptides were similar and *F2A adds an extra 22 aa to the C-terminus of the target protein compared to F2A. However, we noted that the abundance of target and reference proteins appeared to be higher in *F2A samples than F2A samples (Figure 2D). A non-specific protein bound by the GFP antibody had approximately equal abundance in F2A and *F2A samples. In contrast, target and reference proteins were roughly 2-fold higher in the *F2A samples than in F2A samples (Figure 2D, F). This suggested that *F2A promotes more efficient translation of a polycistronic transcript than F2A, and therefore may be a better choice for some applications.

### Comparison of fluorescent and luminescent reporter proteins in tobacco leaves

We set out to identify reporter proteins that would be most detectable after transient expression in tobacco leaves. Various FPs with different spectral properties have been developed and used to analyze the dynamics of protein localization *in planta*, most commonly in roots. Photosynthetic tissues, however, pose a particular challenge for detecting FPs due to high background autofluorescence, e.g. from chlorophyll. We selected four intrinsically bright, monomeric fluorescent reporters to test: mScarlet-I, mNeonGreen, mCerulean-NLS, and Venus. We synthesized plant codon-optimized forms of mScarlet-I, mNeonGreen, and mCerulean-NLS.

We selected FPs that could be paired as dual reporters with minimal spectral overlap. mScarlet-I is a novel bright monomeric RFP (red fluorescent protein) with a Thr74Ile mutation that results in high photostability, fast maturation (< 40 min), and a high quantum yield (0.54) that is 2.5 times brighter than mCherry (Bindels et al., 2017). mScarlet-I has been used in live cell imaging in Arabidopsis (Kimata et al., 2019). In the green range, we chose mNeonGreen, a bright and stable green-yellow fluorescent protein derived from monomerization of the tetrameric yellow fluorescent protein LanYFP (Shaner et al., 2013). mNeonGreen is about 3-5 times brighter than GFP and EGFP, and its maturation time is about 3-fold less than EGFP (Shaner et al., 2013; Cranfill et al., 2016; Rodriguez et al., 2017; Steiert et al., 2018). Despite being a relatively new fluorescent protein, it has been successfully expressed in several plant species such as *A*.*thaliana, N*.*benthamiana*, and rice (Kimata et al., 2019; Kato et al., 2019; Pasin et al., 2014; Stoddard and Rolland, 2019; Luginbuehl et al., 2019). mCerulean is a notable *Aequorea* GFP variant that is cyan in color (Rizzo et al., 2004). It is reported to be a very rapidly maturing monomer. The S72A/Y145A/H148D/A206K amino acid substitutions found in mCerulean make it more photostable and 2.5 times brighter than ECFP (Rizzo et al., 2004; Rizzo and Piston, 2005). In the yellow range, we selected Venus, an improved version of YFP (yellow fluorescent protein) with a novel F46L mutation (Nagai et al., 2002). Due to its enhanced brightness and fast maturation time (<20 min), Venus has been utilized in the development of ratiometric sensors such as DII-VENUS and Jas9-VENUS, which monitor proteins that undergo rapid turnover (Brunoud et al., 2012; Larrieu et al., 2015).

We used spectral scanning to identify excitation and emission wavelengths for each FP that produced the strongest signal above background autofluorescence of tobacco leaves. In some cases, these differed from the peak wavelengths identified from *in vitro* studies of the FPs (Table S2). We performed tests with a KAI2 target protein to evaluate the performance of pRATIO vectors containing mScarlet-I/Venus, mNeonGreen/mScarlet-I, and mScarlet-I/mCerulean reporter pairs (Figure 3A). The constructs were introduced into tobacco leaves by *Agrobacterium*-mediated transformation. After 3 days, fluorescence signals from leaf discs were measured in a microplate reader equipped with linear variable filters (Table S2). Among the four fluorescent proteins, the ratio of signal to background was highest for mScarlet-I, ranging from 40-fold to 156-fold (Figure 3B). The superior performance of mScarlet-I is likely a combination of its exceptional brightness for a red FP and the comparably low autofluorescence from tobacco leaves at its optimal excitation and emission settings (Bindels et al., 2017; Thorn, 2017). Venus also worked well, with a signal intensity 46-fold higher than background. In contrast, background autofluorescence was much higher under mNeonGreen and mCerulean filter settings, limiting the potential dynamic range and utility of these FPs in leaf tissue assays.

**Figure 3.**
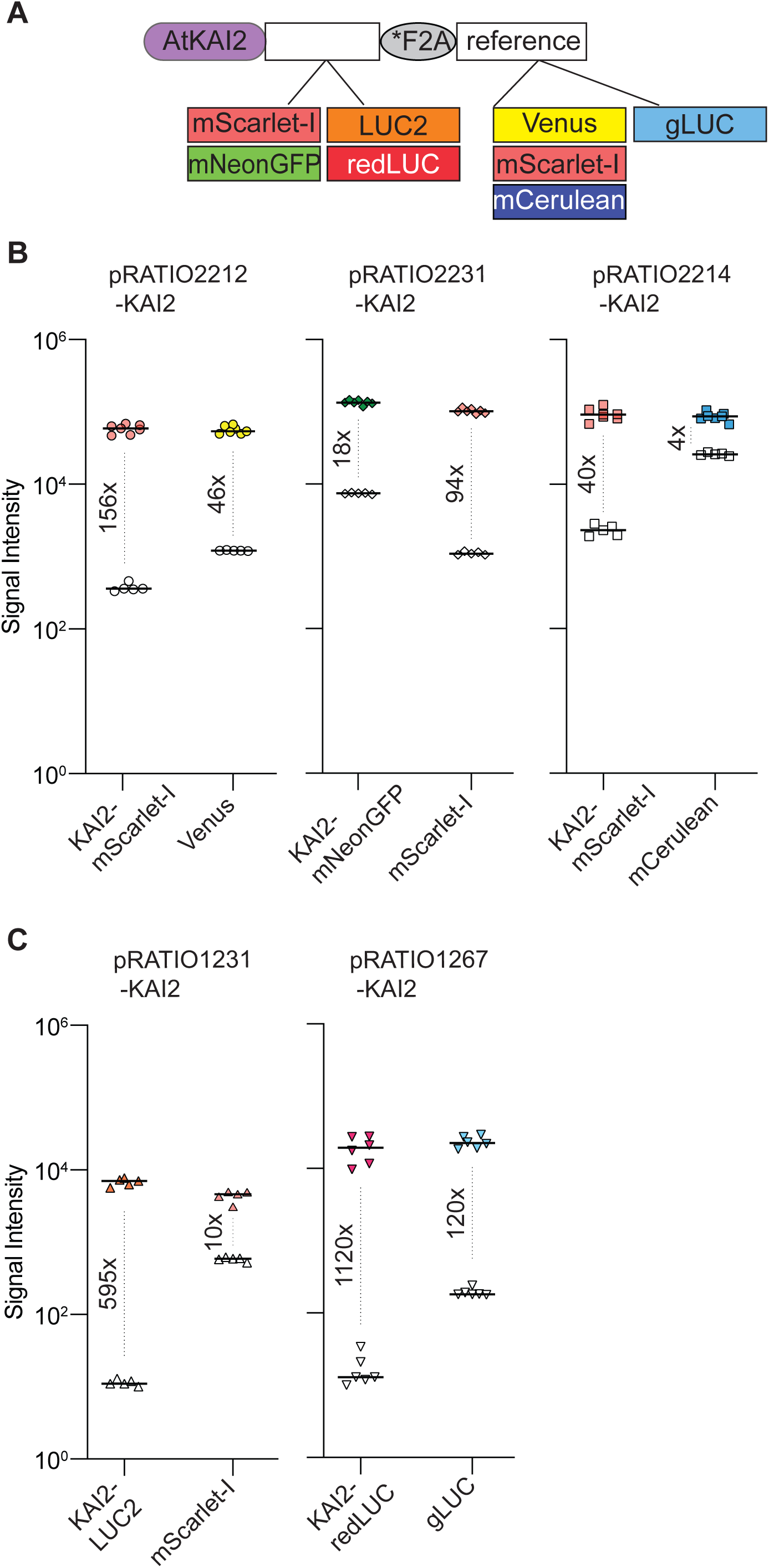
Comparison of signal intensity among pRATIO vectors with different combination of target and reference proteins. **(A)** Schematic of the pRATIO vector expressing Arabidopsis KAI2 cDNA (AtKAI2) fused to different target and reference proteins, **(B, C)** The scatter dot plots comparing signal intensities of the reporter proteins. Leaves transformed with p19 served as the negative control for background signal (open symbols). Fluorescence and luminescence signals were measured in tobacco leaf epidermal cells 72 hpi. Median is shown, n = 4-6 leaf discs. The numbers indicate fold increase in signal intensity over background.

Bioluminescent reporter proteins potentially offer greater sensitivity and dynamic ranges for quantitation than fluorescent reporters, with the drawback that they must be supplied with substrates to enable detection. We selected three luciferases to test. LUC2 is an improved version of the firefly luciferase isolated from *Photinus pyralis (Mašek et al., 2013)*. LUC2 requires ATP and molecular oxygen to catalyze the yellow light-emitting reaction with its substrate, D-luciferin (Marques and Esteves da Silva, 2009). *P. pyralis* luciferase and optimized variants of it have been used in many *in vivo* imaging experiments in plants, perhaps most famously for tracking circadian clock-regulated gene expression over the course of several days. A mutant form of luciferase from the Japanese firefly, *Luciola cruciata*, that has a red-shifted emission spectrum (here referred to as redLUC) was also chosen (Kajiyama and Nakano, 1991; Tafreshi et al., 2008). redLUC also uses D-luciferin as a substrate (Branchini et al., 2005). Because of spectral overlap, however, redLUC and LUC2 are not a suitable pair for dual-luciferase assays. Instead, redLUC combines well in dual-luciferase assays with Gaussia Dura luciferase (gLUC), a mutated blue variant of a luciferase from the marine copepod *Gaussia princeps* that confers stabilized luminescence (Welsh et al., 2009; Markova et al., 2019). gLUC is one of the smallest and brightest luciferases currently known. It catalyzes the oxidative decarboxylation of coelenterazine in an ATP-independent manner to produce blue light with a peak wavelength around 480 nm (Tannous et al., 2005). Because the luminescence from redLUC and gLUC can be spectrally resolved, simultaneous measurement of both reporters can be accomplished without the need for two-step addition of substrates or quenching.

We synthesized coding sequences for *LUC2, redLUC*, and *gLUC* that were codon-optimized for expression in *Arabidopsis thaliana* and generated several pRATIO vectors that incorporate these reporters. We tested pRATIO1231-KAI2 and pRATIO1267-KAI2, which respectively use LUC2/mScarlet-I and redLUC/gLUC as target/reference reporters. In comparison to FPs, firefly luciferase proteins expressed in tobacco leaves had a substantially higher ratio of luminescence signal to background, due to substantially lower background signals. gLUC also performed well, but had higher background signal, possibly due to luciferase-independent decomposition of the coelenterazine substrate (Figure 3C).

### Ratiometric analysis of KAI2 degradation in *N*.***benthamiana***

Having developed ratiometric dual-fluorescent and dual-luminescent reporters, we investigated whether KAR-induced degradation of Arabidopsis KAI2 can be observed in tobacco. We cloned *KAI2* and the catalytically inactive *S95A* allele of *kai2* into pRATIO4212, which uses a *UBQ10* promoter and mScarlet-I/Venus reporters. Transient expression of these constructs in tobacco leaf epidermal cells showed that KAI2-mScarlet-I and kai2^S95A^-mScarlet-I were localized to the cytoplasm and the nucleus (Figure 4A). This was consistent with the subcellular localization of KAI2 in *Arabidopsis* (Sun and Ni, 2011). After 12 h of treatment with 10 µM KAR_2_, we did not observe a decline in the target to reference ratio for either KAI2 or kai2^S95A^ (Figure 4C). We performed similar tests with pRATIO3267, which uses a *35S* promoter and redLUC/gLUC reporters. Again, we did not observe a decline in the target to reference ratio after KAR_2_ treatment with either KAI2 protein (Figure 4D).

**Figure 4.**
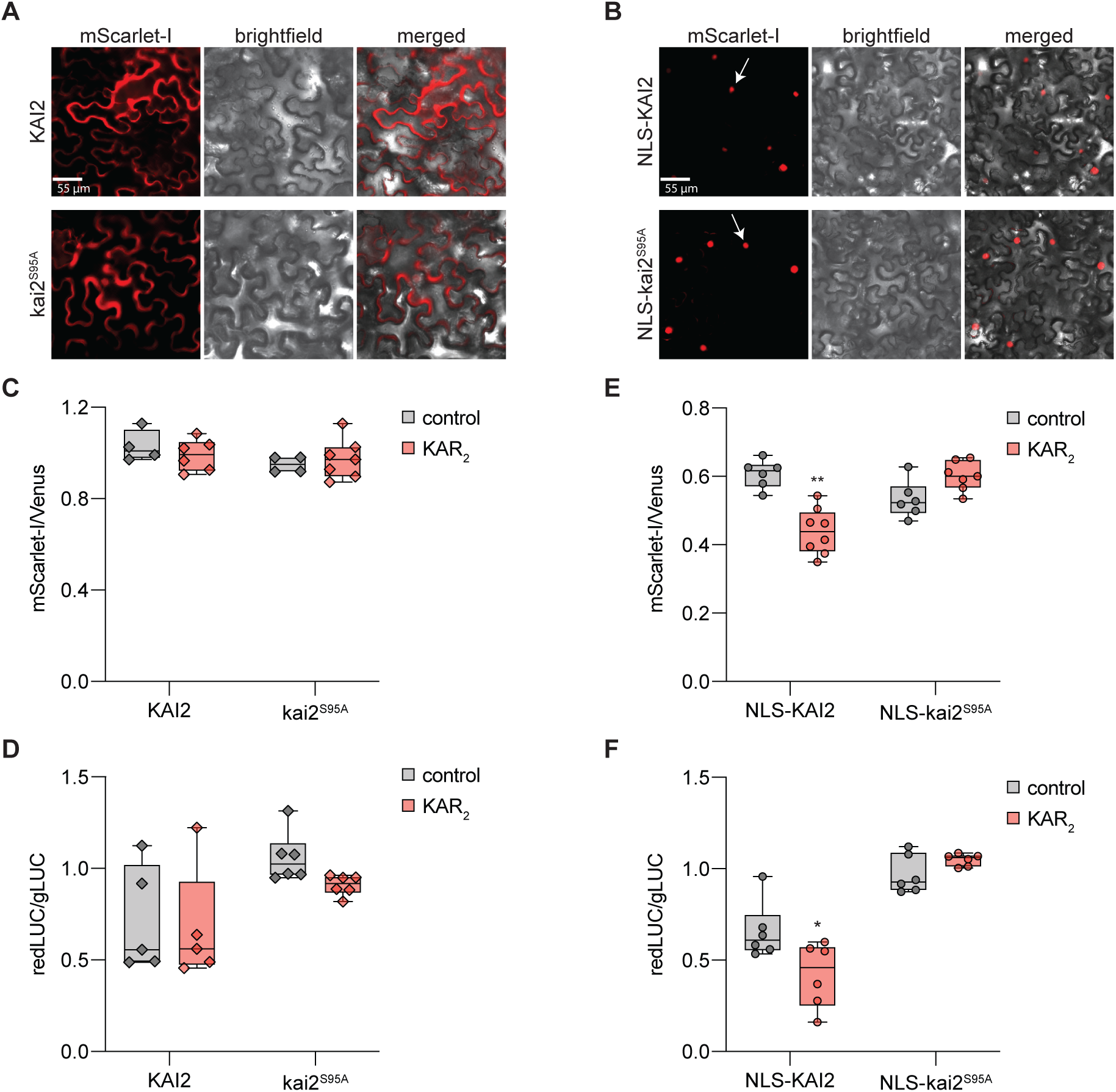
Ratiometric detection of KAI2 degradation in tobacco. **(A, B)** *N*.*benthamiana* leaf epidermal cells expressing wild type *Arabidopsis* KAI2 (AtKAI2) and catalytically inactive mutant of KAI2 (kai2^S95A^) fused to mScarlet-I in **(A)** pRATIO4212 and **(B)** pRATIO2212. Images were visualized using the RFP epifluorescence settings. Arrows indicate nuclear localization. Scale bar = 55 µm. **(C-F)** Box and whisker plots showing KAR_2_ induced degradation response of KAI2 and kai2^S95A^ in transiently transformed tobacco epidermal cells. Degradation response was monitored in **(C, D)** pRATIO4212 and pRATIO3267 (without NLS fusion), **(E, F)** pRATIO2212 and pRATIO1267 (KAI2/*kai2*^*S95A*^ cDNA fused with nuclear localization signal (NLS) at N-terminus in the presence of 10 µM KAR_2_ or 0.02% acetone control. mScarlet-I-to-Venus (mScarlet-I/Venus) and redLUC-to-gLUC (redLUC/gLUC) ratios are plotted on the y-axis. n = 4-7 leaf discs. * *P* < 0.01; ** *P* < 0.001, Mann-Whitney U test comparisons to control treatment.

Unlike KAI2, its signaling partner MAX2 is only found in the nucleus of *Arabidopsis* cells (Stirnberg et al., 2007; Shen et al., 2007). In addition, SMAX1, the target of KAI2, co-localizes with TOPLESS(TPL)/TPL-RELATED(TPR) proteins in the nucleus, which is consistent with the nuclear localization pattern of the homologous D53-type SMAX1-LIKE (SMXL) proteins (Jiang et al., 2013; Zhou et al., 2013; Soundappan et al., 2015; Wang et al., 2015; Liang et al., 2016). This led us to test whether the subcellular localization of KAI2 influences its potential to be degraded after KAR treatment.

Therefore, we examined KAI2 degradation with pRATIO2212 and pRATIO1267, which are respectively identical to pRATIO4212 and pRATIO3267 but add an NLS to the N-terminus of the target protein. Transient expression of pRATIO4212-KAI2 and -kai2^S95A^ in tobacco produced KAI2 fusion proteins that were exclusively localized to the nucleus. In contrast to our prior results, we observed that the target to reference ratio of NLS-KAI2 decreased after KAR_2_ treatment when either the dual-fluorescent or dual-luminescent reporter system was used. No decline was observed for NLS-kai2^S95A^ targets after KAR_2_ treatment (Figure 4E,F). This is consistent with the importance of Ser95 for KAR signaling and the stability of kai2^S95A^ in *Arabidopsis* after KAR treatment (Waters et al., 2015). Our results suggest that nuclear localization is important for KAR_2_-induced degradation of KAI2.

## DISCUSSION

### Functional evaluation of the pRATIO vectors

We developed a set of pRATIO vectors to aid studies of post-translational regulation of proteins (Figure 1). These vectors enable *in vivo* monitoring of dynamics in protein abundance in response to applied stimuli. These assays can be carried out rapidly in transient expression systems. Importantly, pRATIO vectors can normalize differences in transformation efficiency or transgene expression across samples by simultaneously expressing a reference protein from the same transcript as a target protein of interest. This is made possible by the use of a short “self-cleaving” F2A peptide derived from FMDV. To date, there has not been a comprehensive study that compares the cleavage efficiencies of different 2A peptides in plants. Of the various 2A and 2A-like peptides, the most widely used 2A sequence in plants is F2A (Halpin et al., 1999; El Amrani et al., 2004; Samalova et al., 2006; François et al., 2004; Ma and Mitra, 2002; Burén et al., 2012). We tested two long versions of the F2A peptide, one of which included a putative protease cleavage site that proved to be ineffective. We found that F2A and *F2A have similarly high cleavage efficiencies but that *F2A allows better protein expression (Figure 2). We identified the fluorescent proteins mScarlet-I and Venus, and the bioluminescent proteins redLUC and gLUC, as particularly useful reporter pairs for ratiometric target detection in tobacco leaves (Figure 3). Their superior performance is likely due to the low background signal from green plant tissue at their detection filter settings. The pRATIO vectors have a modular configuration in which the promoter, Gateway cassette, 2A peptide, reporters are flanked by unique restriction endonuclease cleavage sites (Figure 1). This makes further modification of pRATIO vectors to fit specific experimental needs easy to accomplish.

We found that *Arabidopsis* KAI2 was degraded in tobacco following KAR_2_ treatment, demonstrating that this response is conserved between the two species (Figure 4; Waters et al., 2015). KAR-induced degradation of KAI2 was only observed when KAI2 was retained in the nucleus. Interestingly, MAX2 and SMAX1, the signaling partners of KAI2, are nuclear proteins. Although KAI2 degradation is known to be MAX2-independent, future work should investigate whether it depends on KAI2 association with SMAX1 in the nucleus.

### Limitations of the pRATIO system

We note three important limitations when using the pRATIO vectors. First, an appropriate filter set is critical to maximize the signal and reduce spectral overlap between fluorescent proteins (Tables S2, S3). We used spectral scanning to identify optimal excitation and emission settings for each fluorophore in green leaves. However, when typical filter settings for GFP (excitation 488 nm; emission 507 nm), mCherry (excitation 587 nm; emission 610 nm), and CFP (excitation 433 nm; emission 475 nm) were used to detect mNeonGreen, mScarlet-I, and mCerulean, respectively, the fluorescence signal over background was greatly diminished (data not shown). Thus, the use of some of these reporters may be limited by the availability of a multi-mode plate reader that can set continuously adjustable wavelengths and bandwidths for excitation and emission.

Second, although our *F2A sequence produced efficient cleavage and stronger expression of target and reference proteins than F2A, it was not removed from the target post-translationally as anticipated. It is possible that the C-terminal extension may interfere with the function of a target protein or with the activity of reporter proteins. This may not pose a significant problem for pRATIO vectors, as we were able to detect all reporter proteins effectively (Supplemental Figure 1). In cases where this is a problem, however, a better alternative to *F2A may be IntF2A, a fusion of an Ssp DnaE mini-intein variant to a long 58 aa F2A peptide (Zhang et al., 2017). IntF2A is rapidly and efficiently removed from the C-terminal end of a fusion protein through the hyper-N-terminal auto-cleaving action of the intein after translation. IntF2 has been shown to be processed efficiently in multiple organs of transgenic *Nicotiana tabacum*, and in transiently transformed *N. benthamiana* and lettuce (Zhang et al., 2017).

Third, we found pRATIO vectors to be useful in transient expression assays, but we were unable to detect the Venus or mCerulean-NLS reference proteins in transgenic *Arabidopsis thaliana* lines carrying pRATIO2212-KAI2 or pRATIO2214-KAI2 by fluorescence microscopy (Supplemental Figure 3). One likely explanation is that the ribosome does not always continue translation after disruption of the glycyl-prolyl bond at the end of the 2A peptide and instead drops off (Ryan et al., 1999; Donnelly et al., 2001a; de Felipe et al., 2003; Liu et al., 2017). The high level of transient expression that can be achieved in tobacco may overcome an inefficiency in translating the reference protein. Notably, the 2A-dependent ratiometric sensors for auxin and strigolactone have only been deployed in transient expression experiments in protoplasts, raising the question of whether they are effective as stable transgenes (Wend et al., 2013; Samodelov et al., 2016).

Alternatively, 2A peptides may have different effects in different species, depending on how they interact with the ribosome. For example, the cleavage efficiencies of the 2A peptide from FMDV have ranged from 40% to 90% to nearly 100% (Donnelly et al., 2001a; Szymczak et al., 2004b; Kim et al., 2011). This variability is likely due to differences in experimental conditions, including the use of different model organisms. 2A variants found in other viruses, such as equine rhinitis A virus 2A (E2A), porcine teschovirus-1 2A (P2A), thosea asigna virus 2A (T2A), can have different activities in a given system (Ryan et al., 1991; Donnelly et al., 2001a; Szymczak and Vignali, 2005). The 22 aa version of F2A produces the least efficient cleavage of four 2A peptides tested in human cell lines, zebrafish embryos, and mouse liver. P2A is most effective, in some cases producing more than twice the cleavage efficiency of F2A (Kim et al., 2011). It is possible that other 2A forms may be more effective than F2A at inducing stop-and-go translation in transgenic plants. It will be interesting to determine whether there is a tradeoff between cleavage efficiency and the frequency of continued translation of the second coding sequence. We propose that a high cleavage efficiency should be prioritized for accurate monitoring of target/reference ratios.

### Future applications for ratiometric reporters

There are a number of potential applications for controlled co-expression of target and reference proteins from a single polycistronic mRNA, particularly if the reference protein can be detected in stably transformed plants. For example, the *35S* or *UBQ10* promoters in a pRATIO construct could be replaced with a native promoter and coding sequence for a gene of interest. This would potentially enable monitoring of the protein distribution and transcriptional pattern of a gene in a single construct by visualizing the target and reference reporters, respectively. In addition to revealing differences in localization patterns, such a system could be used to simultaneously examine changes in gene expression at the transcript and protein levels. In the case of understanding the nature of KL, we anticipate development of a reliable reporter of KAI2 signaling activity will facilitate KL identification through bioassay-guided fractionation. Alternatively, a ratiometric reporter of KAI2 signaling may enable genetic screens for KL-deficient or -overproducing mutants.

## SUPPLEMENTAL DATA

**Supplemental Figure 1**. Fluorescence microscopy images showing the expression of reference proteins in tobacco epidermal cells.

**Supplemental Figure 2**. Western blot analysis revealing cleavage efficiency in the 2As in tobacco epidermal cells.

**Supplemental Figure 3**. Reference protein is undetectable in *Arabidopsis thaliana*.

**Supplemental Table 1**. List of vectors in the pRATIO series.

**Supplemental Table 2**. Fluorescent proteins (FPs) used in this study.

**Supplemental Table 3**. Luciferases used in this study.

**Supplemental Table 4**. Primers used in this study.

**Supplemental Table 5**. GenBank accession numbers for the pRATIO vectors.

## ACKNOWLEDGEMENTS

We gratefully acknowledge funding support from NSF IOS-1350561 (now IOS-1737153) and NSF IOS-1557962 (now IOS-1740560) to DCN, and NSF ISO-1751385 to JMVN. We thank Dr. Gavin Flematti and Dr. Adrian Scaffidi (University of Western Australia) for supplying KARs.

## AUTHOR CONTRIBUTIONS

Project and experimental design by AK and DCN. Experiments were carried out by AK, SK, and CR. All authors contributed to data analysis and interpretation. Figure preparation by AK and DCN. Manuscript preparation by AK, JMVN, and DCN.

## METHODS

### Construction of plant transformation vectors

#### Biological components

The plant codon optimized coding sequence of *mScarlet-I, LUC2, mCeruelan-NLS*, and *redLUC-*F2A-gLUC* were synthesized in pUC57 cloning vectors (Genscript) using *KpnI - SacI, ApaI-MluI, KpnI-SacI*, and *MluI-SacI* restriction sites, respectively. The *Arabidopsis UBQ10* promoter was derived from pUBQ10:YFP-GW plasmid (Michniewicz et al., 2015), plant codon optimized *mNeonGreen* from pMCS:mNeonGreen-GW plasmid (Lucia Strader, Washington University), and NLS from SV40 large T antigen.

pUC57-mScarlet-I: mScarlet-I in pUC57

pUC57-mCerulean: mCerulean-NLS in pUC57

pUC57-LUC2: LUC2 in pUC57

pUC57-redgLUC: redLUC-P-2A-gLUC in pUC57

pUC57-mNeonGreen: pUBQ10-NLS-GW-mNeonGreen-F2A-mCherry in pUC57

pUC57-mNeonGreen was used as a template to generate pRATIO2131. mScarlet-I was excised from pUC57-mScarlet-I using *KpnI*-*SacI* and inserted into the corresponding site of pUC57-mNeonGreen to generate pUC2131. The *HindIII-SacI* fragment of the resulting plasmid was ligated into the same sites of pGWB401 (Nakagawa et al., 2007) to generate pRATIO2131.

To create pRATIO2112, the coding sequence of *Venus* was amplified from pCN-SANB-nu3V (Wolfgang Busch, Salk Institute) using primers that introduce 5’ *KpnI* site and a 3’ *SacI* site. The PCR product was digested with *KpnI-SacI* and inserted into *KpnI-SacI* digested pUC57-mNeonGreen. The coding sequence of *mScarlet-I* was amplified from pUC57-mScarlet-I with gene specific primers introducing a 5’ *ApaI* site and 3’ *Mlu*I site. The resulting PCR product was digested with *ApaI-MluI* and inserted into the corresponding site of pUC57-mNeonGreen to generate pUC2112. The resulting pUBQ10-NLS-GW-mScarlet-I-F2A-Venus was released from pUC57 using *HindIII-SacI* and inserted into *HindIII-SacI* digested pGWB401. Oligonucleotides used for PCR amplifications are listed in Table S4.

To generate pRATIO2151, LUC2 was excised from pUC57-LUC2 using *ApaI*-*MluI* and cloned into *ApaI-MluI* digested pUC2131 to generate pUC2151. The *HindIII-SacI* fragment of the resulting plasmid was ligated into the same sites of pGWB401 to generate pRATIO2151.

To create pRATIO1112, pUBQ10-NLS-GW-mScarlet-I-F2A-Venus was released from pUC2112 using *XbaI* and *SacI* restriction enzymes and cloned into *XbaI - SacI* digested pGBW402 (Nakagawa et al., 2007). pRATIO1131 and 1151 were made in a similar fashion using pUC2131 and pUC2151 as template, respectively.

*F2A was excised from pUC57-redgLUC using *MluI* and *KpnI*, and subcloned into the same sites of pRATIO1112 and 1151 to create pRATIO1212 and 1251, respectively. To generate pRATIO1267, pUC57-redgLUC was digested with *ApaI* and *SacI* and inserted into the corresponding site of pRATIO1212.

To construct pRATIO2212, pRATIO2251, and pRATIO2231, *F2A was released from pUC57-redgLUC using *MluI* and *KpnI* and ligated into the corresponding sites of pRATIO2112, 2151, and 2231, respectively.

To create pRATIO2214, mCerulean-NLS was excised from pUC57-mCerulean using *KpnI* and *SacI* and ligated into *KpnI-SacI* digested pRATIO2212.

To construct the no NLS versions of pRATIO, NLS sequence was removed by digesting pRATIO1212, pRATIO2212-2231 with *XbaI-SpeI*, followed by self-ligation to generate pRATIO3212, pRATIO4212-4231, respectively.

### Transient expression in *N*.***benthamiana***

*Nicotiana benthamiana* were grown in soil in a growth room at 22°C under long day conditions (16/8 hr light dark cycle). *N. benthamiana* leaves (3 weeks old) were infiltrated with *A. tumefaciens* strain GV3101 harboring pRATIO-KAI2 fusion vectors. GV3101 cells were grown in 10 ml LB broth with antibiotics overnight at 28°C and then pelleted at 2,500 x*g* for 10 min. Cells were then washed in 10 ml of infiltration medium (10 mM MES pH 5.7, 10 mM MgCl_2_ and 150 μM acetosyringone), centrifuged again at 2,500 x*g* for 5 min and resuspended in infiltration medium at an OD_600_ of 0.2. The *Agrobacterium* solution was infiltrated in the abaxial surface of the leaf using a 1 ml syringe without a needle. After infiltration plants were placed under normal light conditions and leaves were collected after 72 hours for further analysis.

### Degradation assays in tobacco

To generate pRATIO vectors with KAI2 and kai2^S95A^ fusions, full length *KAI2* coding sequence was amplified from Col-0 cDNA and inserted into Gateway entry vector pDONR207. The resulting entry clone was then moved to the pRATIO destination vectors by gateway LR reaction. The *kai2*^*S95A*^ mutant was generated with Infusion on *KAI2* entry clone described above, and subsequently recombined with pRATIO vectors.

To perform degradation assays, leaf discs were excised 3 days post infiltration and incubated at RT for 12 hr in the presence or absence of 10 µM KAR_2_ or 0.02% acetone control. mScarlet-I (target) and Venus (reference) were excited at 560 ± 10 nm and 497 ± 15 nm in a CLARIOstar plate reader (BMG Labtech, Ortenberg, Germany). Emission was recorded at 595 ± 10 nm (mScarlet-I) and 540 ± 20 nm (Venus) using black 96-well plates (Costar). Degradation was quantified as mScarlet-I/Venus fluorescence intensity ratios after background subtraction using p19 transformed leaf disc.

For luminescence-based degradation assay, redLUC and gLUC were detected at 640 ± 10 nm and 480 ± 40 nm, respectively using white 96-well plates (PerkinElmer).

### Fluorescence and luminescence measurements

Fluorescence and luminescence were measured off-line using the CLARIOstar plate reader (BMG Labtech, Ortenberg, Germany). Fluorescence intensity was measured in well scan mode of density 9×9 (9 points per well) in black, flat bottom plates. For mScarlet-I, Venus, mNeonGreen, and mCerulean detection, the optic settings (excitation, dichroic, and emission) to obtain best signal to noise ratio are listed in Table S2. Luminescence was measured in endpoint mode in white, flat bottom plates (costar). Table S3 shows the optimal filters to detect LUC2, redLUC, and gLUC signals in *N*.*benthamiana*.

Tobacco leaf discs were imaged 3 days after agroinfiltration on Keyence BZ-X710 epi fluorescence microscope. Fluorescence was observed using the RFP filter setting (Ex 470/40 nm, Em 535/50nm) to detect mScarlet-I.

### Protein extraction and western blot analysis

Transformed tobacco leaves were ground in liquid nitrogen and then resuspended hot SDS-sample buffer (100 mM Tris-HCl, pH 6.8, 2% SDS, 20% glycerol, 100 mM DTT, and 0.004% bromophenol blue, 0.48 g Urea per ml). Samples were then cleaned by centrifugation at 10,000 xg at room temperature. For western blotting analysis, samples were separated on SDS-PAGE and the proteins were transferred onto PVDF membrane. Subsequently, the membrane was probed with the indicated primary antibody (rabbit anti-GFP [1:1500, Abcam, #ab290], mouse anti-RFP [1:1000, Chromotek, #6G6], and mouse anti-2A [1:1000, Sigma, #3H4]), washed with TBST, and probed with HRP-conjugated mouse anti-rabbit (1:10,000, Genscript, #A01856) and horse anti-mouse (1:5000, Cell Signaling, #7076). Blots were developed using an Azure Radiance Plus chemiluminescent substrate (#AC2102).

