## Supplemental files for "A series of dual-reporter vectors for ratiometric analysis of protein abundance in plants"

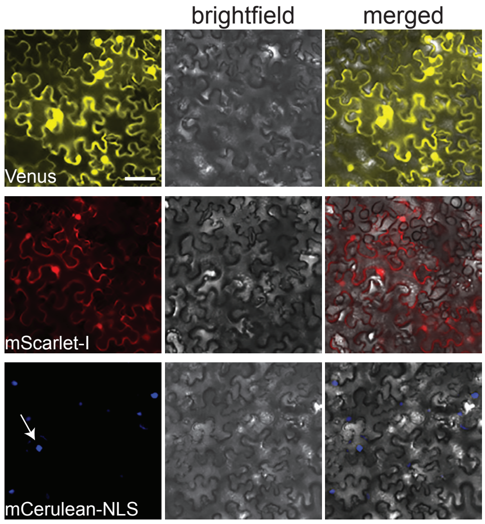


**Supplemental Figure 1.** Fluorescence microscopy images showing the expression of reference proteins in tobacco epidermal cells.

Epifluorescence microscopy images of *N.benthamiana* epidermal cells transformed with pRATIO2212, 2231, and 2214, expressing Venus (yellow), mScarlet-I (red), and mCeruelan-NLS (blue) as reference proteins, respectively. Arrow indicates nuclear localization. Scale bar = 55 µm.

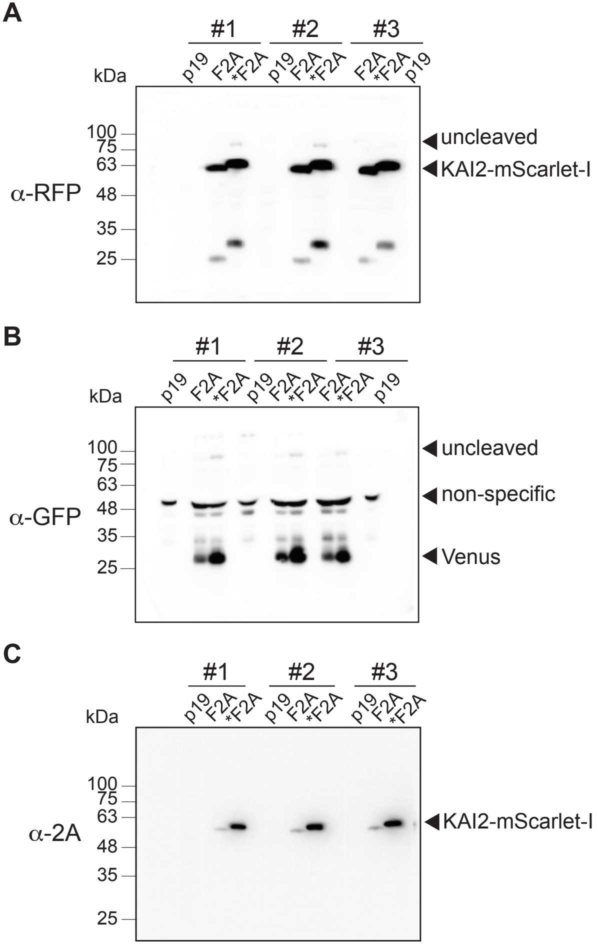


**Supplemental Figure 2. Western blot analysis revealing cleavage efficiency in the 2As in tobacco epidermal cells.**

Proteins obtained from three independent biological replicates of each F2A and *F2A samples were probed against **(A)** mScarlet-I, **(B)** Venus, and **(C)** 2A peptide. The cleavage efficiency was assessed as shown in Figure 2D. Leaf transformed with p19 served as the negative control.

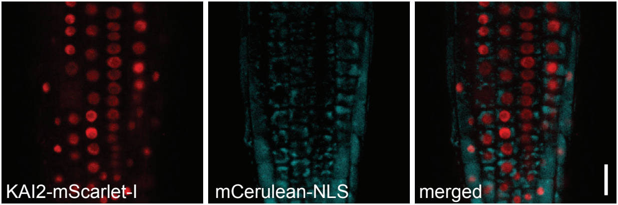


**Supplemental Figure 3.** Reference protein is undetectable in *Arabidopsis thaliana*.

Confocal microscopy of *Arabidopsis* root meristem in homozygous transgenic lines expressing KAI2-mScarlet-I fusion protein from the *UBQ10* promoter in pRATIO2214. Scale bar: 50 µm

**Table S1.** List of vectors in the pRATIO series.

| pRATIO | Promoter | NLS (+/-) | Target | 2A | Reference | Selection marker (Bacteria/Plants) |
|---|---|---|---|---|---|---|
| **pRATIO1112** | *35Sp* | + | mScarlet-I | F2A | Venus | Spec/Kan |
| **pRATIO1131** | *35Sp* | + | mNeonGFP | F2A | mScarlet-I | Spec/Kan |
| **pRATIO1151** | *35Sp* | + | LUC2 | F2A | mScarlet-I | Spec/Kan |
| **pRATIO2112** | *UBQ10p* | + | mScarlet-I | F2A | Venus | Spec/Kan |
| **pRATIO2131** | *UBQ10p* | + | mNeonGFP | F2A | mScarlet-I | Spec/Kan |
| **pRATIO2151** | *UBQ10p* | + | LUC2 | F2A | mScarlet-I | Spec/Kan |
| **pRATIO1212** | *35Sp* | + | mScarlet-I | *F2A | Venus | Spec/Kan |
| **pRATIO1251** | *35Sp* | + | LUC2 | *F2A | mScarlet-I | Spec/Kan |
| **pRATIO1267** | *35Sp* | + | redLUC | *F2A | gLUC | Spec/Kan |
| **pRATIO2212** | *UBQ10p* | + | mScarlet-I | *F2A | Venus | Spec/Kan |
| **pRATIO2214** | *UBQ10p* | + | mScarlet-I | *F2A | mCerulean | Spec/Kan |
| **pRATIO2231** | *UBQ10p* | + | mNeonGFP | *F2A | mScarlet-I | Spec/Kan |
| **pRATIO2251** | *UBQ10p* | + | LUC2 | *F2A | mScarlet-I | Spec/Kan |
| **pRATIO3212** | *35Sp* | - | mScarlet-I | *F2A | Venus | Spec/Kan |
| **pRATIO3267** | *35Sp* | - | redLUC | *F2A | gLUC | Spec/Kan |
| **pRATIO4212** | *UBQ10p* | - | mScarlet-I | *F2A | Venus | Spec/Kan |
| **pRATIO4214** | *UBQ10p* | - | mScarlet-I | *F2A | mCerulean | Spec/Kan |
| **pRATIO4231** | *UBQ10p* | - | mNeonGFP | *F2A | mScarlet-I | Spec/Kan |

**Table S2.** Fluorescent proteins (FPs) used in this study.

| | Optimal filter settings for tobacco | | | Type | Brightness | Photostability | Maturation[e] |
|---|---|---|---|---|---|---|---|
| FP | Excitation | Dichroic | Emission | | E X QY[f] | $t_{1/2}$ (s)[g] | $t_{50}$ (min)[h] |
| mScarlet-I[a] | 560-10 | 573.5 | 595-10 | m | 57 | 225 | 36 |
| Venus[b] | 497-15 | 517.2 | 540-20 | m | 53 | 15 | 17.6 |
| mNeonGreen[c] | 505-10 | 522.5 | 540-10 | m | 92.8 | 158 | 10 |
| mCerulean[d] | 420-10 | 446.5 | 473-10 | m | 17 | NA | 6.6 |

[a]Bindels et al. 2017, [b]Nagai et al. 2002, [c]Shaner et al. 2013, [d]Rizzo et al. 2004, [e]Balleza et al. 2018, [f] E is the extinction coefficient, QY is the quantum yield, and calculated brightness is the product of E and QY (E × QY) is their product. [g] time in seconds to bleach to half of the initial intensity at an initial emission rate of 1000 photons/s. [h]Time of half-maximal fluorescence maturation ($t_{50}$). NA, not determined. Type: m, monomer.

**Table S3.** Luciferases used in this study.

| Luciferase | Origin | Optimal filter settings | Substrate |
|---|---|---|---|
| LUC2 | *Photinus pyralis* | 580-80 | D-Luciferin |
| redLUC | *Luciola cruciata* | 640-20 | D-Luciferin |
| gLUC | *Gaussia princeps* | 480-80 | Coelenterazine |

**Table S4.** Primers used in this study.

| Name | 5'-3' sequence |
|---|---|
| mScarlet_ApaI-F | CAT TCG CG**G GGC CCA** ATG GTG TC |
| mScarlet_MluI-R | CAGTGAATTCGAG**ACGCGT**CTTGTACAACT |
| Venus_KpnI-F | GTACAAAGTG**GGTACC**ATGGTGAGCA |
| Venus_SacI-R | CTT**GAGCTC**TTAGGCCGCTGCAGCAAT |
| KAI2_GW-F | GGGGACAAGTTTGTACAAAAAAGCAGGCTTCATGGGTGTGGTAGAAG |
| KAI2_GW-R | GGGGACCACTTTGTACAAGAAAGCTGGGTGTCACATAGCAATGTCATTAC |
| S95A_Inf-F | GGCCACGCTGTTTCTGCCATGATT |
| S95A_Inf-R | AGAAACAGCGTGGCCAACAAA |

Nucleotide bases shown in bold denote restriction sites used for cloning

**Table S5.** GenBank accession numbers for the pRATIO vectors.

| Vector name | GenBank accession No. |
|---|---|
| pRATIO1112 | MT024582 |
| pRATIO1131 | MT024581 |
| pRATIO1151 | MT024579 |
| pRATIO2112 | MT024574 |
| pRATIO2131 | MT024573 |
| pRATIO2151 | MT024572 |
| pRATIO1212 | MT024583 |
| pRATIO1251 | MT024580 |
| pRATIO1267 | MT024584 |
| pRATIO2212 | MT024578 |
| pRATIO2214 | MT024575 |
| pRATIO2231 | MT024577 |
| pRATIO2251 | MT024576 |
| pRATIO3212 | MT024588 |
| pRATIO3267 | MT024589 |
| pRATIO4212 | MT024587 |
| pRATIO4214 | MT024585 |
| pRATIO4231 | MT024586 |
